# Machine Learning Identifies Genes Linked to Neurological Disorders Induced by Equine Encephalitis viruses (EEV), Traumatic Brain Injuries (TBI), and Organophosphorus nerve agents (OPNA)

**DOI:** 10.1101/2024.11.21.624720

**Authors:** Liduo Yin, Morgen VanderGiessen, Vinoth Kumar, Benjamin Conacher, Po-Chien Haku Chao, Michelle Theus, Erik Johnson, Kylene Kehn-Hall, Xiaowei Wu, Hehuang Xie

**Affiliations:** Department of Biomedical Sciences and Pathobiology, Virginia-Maryland College of Veterinary Medicine, Virginia Polytechnic Institute and State University, Blacksburg, VA 24061, USA; Center for Emerging, Zoonotic, and Arthropod-borne Pathogens, Virginia Polytechnic Institute and State University, Blacksburg, VA 24061, USA; Neuroscience Department, Medical Toxicology Division, U.S. Army Medical Research Institute of Chemical Defense, Aberdeen Proving Ground, MD 21010, USA; Department of Statistics, Virginia Polytechnic Institute and State University, Blacksburg, VA 24061, USA

## Abstract

Venezuelan, eastern, and western equine encephalitis viruses (collectively referred to as equine encephalitis viruses---EEV) cause serious neurological diseases and are a significant threat to the civilian population and the warfighter. Likewise, organophosphorus nerve agents (OPNA) are highly toxic chemicals that pose serious health threats of neurological deficits to both military and civilian personnel around the world. Consequently, only a select few approved research groups are permitted to study these dangerous chemical and biological warfare agents. This has created a significant gap in our scientific understanding of the mechanisms underlying neurological diseases. Valuable insights may be gleaned by drawing parallels to other extensively researched neuropathologies, such as traumatic brain injuries (TBI). By examining combined gene expression profiles, common and unique molecular characteristics may be discovered, providing new insights into medical countermeasures (MCMs) for TBI, EEV infection and OPNA neuropathologies and sequelae. In this study, we collected transcriptomic datasets for neurological disorders caused by TBI, EEV, and OPNA injury, and implemented a framework to normalize and integrate gene expression datasets derived from various platforms. Effective machine learning approaches were developed to identify critical genes that are associated, either shared among the three neuropathologies or to either TBI, EEV, and OPNA. With the aid of deep neural networks, we were able to extract important association signals for accurate prediction of different neurological disorders by using integrated gene expression datasets of VEEV, OPNA, and TBI samples. Gene ontology and pathway analyses further identified neuropathologic features with specific gene product attributes and functions, shedding light on the fundamental biology of these neurological disorders. Collectively, we highlight a workflow to analyze published transcriptomic data using machine learning, which can be used for both identification of gene biomarkers that are unique to specific neurological conditions, as well as genes shared across multiple neuropathologies. These shared genes could serve as potential neuroprotective drug targets for conditions like EEV, TBI, and OPNA.

## Introduction

Neurological disorders caused by infectious diseases, chemical exposures, and physical trauma pose significant public health challenges and are critical concerns in military medicine worldwide. Venezuelan, eastern, and western equine encephalitis viruses (collectively referred to as EEV in this manuscript), organophosphorus nerve agents (OPNA), and traumatic brain injuries (TBI) are particularly significant and interrelated threats to both civilian populations and military personnel. Despite their distinct etiologies, these conditions share common features in their neurological manifestations and potential for severe long-term consequences [1–4] suggesting possible overlapping molecular mechanisms that could be leveraged for therapeutic development.

EEVs represent a significant threat to both human and animal health throughout the Americas. Venezuelan equine encephalitis virus (VEEV), eastern equine encephalitis virus (EEEV), and western equine encephalitis virus (WEEV) belong to the genus Alphavirus in the family *Togaviridae* [5, 6] and are transmitted primarily through mosquito, but have also been weaponized for use as a potential bioweapon by both the US and Soviet Union. These viruses can cause severe neurological diseases, with mortality rates ranging from 1% (VEEV) to 70% (EEEV) [7–9]. Previous research has demonstrated that these viruses cause systemic infection which is either asymptomatic, or presents as a mild flu-like illness in the acute phase of infection [10]. Neuroinvasion occurs around day 4 post infection through the olfactory epithelium, but can also increase blood brain barrier (BBB) permeability to enter the brain via transcytosis, but this is less thoroughly understood for EEEV and WEEV [11, 12]. Recent studies using animal models have revealed that VEEV infection triggers a cascade of inflammatory responses, including the activation of pro-inflammatory cytokines such as IL-1β, TNF-α, and IFN-γ, which contribute to neuronal damage and subsequent neurological symptoms [13]. Despite significant advances in understanding their pathogenesis, current therapeutic options are limited to supportive care, with no specific antiviral treatments available.

Organophosphorus nerve agents (OPNAs) are highly toxic chemicals that interfere with the normal functioning of the nervous system. These compounds, including G-series agents (tabun, sarin, soman) and V-series nerve agents (VE, VG, VM, VR, VX), irreversibly inhibit acetylcholinesterase, leading to excessive accumulation of acetylcholine at synapses [14]. Research over the past decades has revealed that OPNA toxicity extends beyond acute cholinergic crisis. Studies have shown that OPNA exposure initiates complex cellular cascades involving oxidative stress, neuroinflammation, and excitotoxicity [15]. Long-term studies in animal models and human survivors have documented persistent neurological deficits, including cognitive impairment, anxiety, and depression [16, 17]. Current treatment protocols rely primarily on a combination of anticholinergic drugs (such as atropine), oximes for enzyme reactivation, and anticonvulsants[18–21]. However, these treatments must be administered rapidly after exposure and may not prevent long-term neurological consequences. Recent research has focused on understanding the molecular mechanisms of delayed neurotoxicity and developing more effective neuroprotective strategies.

Traumatic brain injury (TBI) is a major global health concern, affecting an estimated 69 million people worldwide each year [22]. The spectrum of TBI ranges from mild concussions to severe injuries with devastating consequences. The pathophysiology of TBI involves both primary injury mechanisms (direct mechanical damage) and secondary injury (inflammatory cascades, BBB breakdown, hemorrhage) that can persist for months or years after the initial trauma [23]. Extensive research has identified key molecular pathways involved in TBI pathogenesis, including neuroinflammation, oxidative stress, excitotoxicity, and disruption of the blood-brain barrier. Recent studies have revealed the complexity of TBI’s molecular signature, with altered expression of numerous genes involved in inflammation (e.g., IL-1β, TNF-α), cell death pathways (e.g., caspase-3, BAX), and neuroplasticity (e.g., BDNF, NGF) [24, 25]. Advanced neuroimaging techniques combined with molecular studies have demonstrated that TBI triggers both acute and chronic changes in brain structure and function [26]. Despite this growing understanding, therapeutic options remain limited, with most treatments focusing on symptom management rather than addressing the underlying pathological mechanisms.

Research on the pathogenesis of these conditions faces numerous challenges. For EEVs, the requirement for high-containment facilities and the complexity of viral-host interactions have limited comprehensive studies [27]. OPNA research faces similar chemical safety concerns, along with ethical considerations that restrict human studies. While TBI research has progressed more rapidly due to greater accessibility and established animal models [28], many aspects of its molecular pathology remain poorly understood. However, recent advances in high-throughput genomic technologies and bioinformatics approaches have opened new avenues for investigating these conditions through comparative analysis of gene expression profiles. This approach allows us to leverage the extensive body of research in one area to inform the understanding of related conditions, potentially identifying common pathways and novel therapeutic targets. By implementing machine learning techniques, we can now integrate and analyze complex transcriptomic datasets from various experimental platforms, identifying both shared and condition-specific molecular signatures [29, 30]. This integrated approach not only provides insights into the fundamental biology of these neurological disorders but also has the potential to guide the development of medical countermeasures that could be effective across multiple conditions. Understanding the commonalities and differences in gene expression patterns among these disorders may reveal new therapeutic targets, ultimately leading to more effective interventions for affected individuals.

This study aims to systematically analyze and compare the transcriptomic profiles associated with TBI, EEV infection, and OPNA exposure, using advanced computational methods to address the challenges of integrating data from diverse experimental platforms and conditions. By utilizing artificial neural network, we identified critical genes and pathways that may provide clues for new therapeutic targets or diagnostic markers. Our findings may contribute to the development of more effective treatments for these neurological conditions.

## Results

### Workflow implemented in this study

In this study, we endeavored to extract salient association signals from integrated gene expression datasets derived from VEEV, OPNA, and TBI samples, with the objective of enabling accurate prediction of diverse neurological disorders. Towards this end, we acquired and reanalyzed a compendium of 6 datasets encompassing 395 samples pertaining to VEEV, OPNA, and TBI, across various experimental conditions and organismal models. These datasets underwent a rigorous normalization and integration process to facilitate downstream analysis. Simultaneous differential expression analysis was performed on each condition to identify informative key genes exhibiting substantial expression variations relative to control samples. Subsequently, the expression matrix of these selected key genes was extracted from the integrated dataset and utilized as input for machine learning algorithms, enabling the identification of distinctive features associated with different neurological diseases (**Figure 1**).

**Figure 1.**
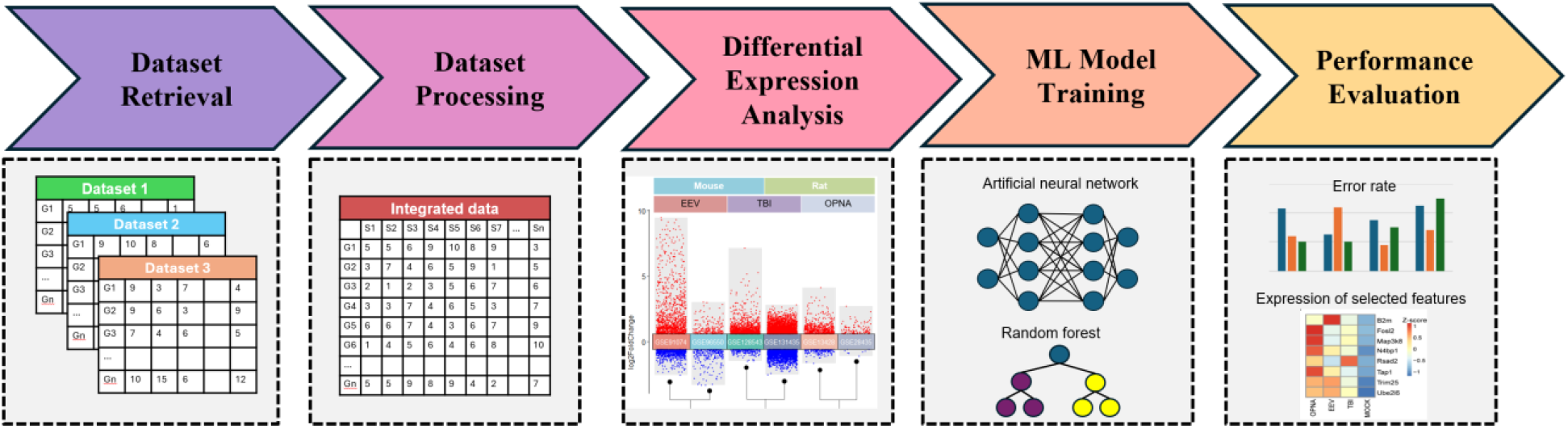
Workflow implemented in this study.

### Collection, normalization and integration of gene expression datasets for OPNA, EEV and TBI derived from various platforms

To investigate the transcriptional characteristics of rodent brains injured by EEV, TBI or OPNA, we systematically searched and downloaded multiple datasets from NCBI’s Gene Expression Omnibus (GEO) database, ensuring representation of each condition. The acquired data encompassed gene expression profiles for these disorders and their corresponding control samples, which consist of 395 samples across two mammalian species, mouse *(Mus musculus*) and rat *(Rat norvegicus)* (**Figure 2**). As a prerequisite to downstream analysis, data normalization was conducted by identifying unique genes across all platforms used in the study. To create a comprehensive gene set, we compiled a list of all involved genes for each dataset and then obtained the intersection set. This process allowed us to identify orthologous genes and determine which ones were similar across species. To account for batch effects caused by diverse experimental designs across the different GEO datasets, we applied the ComBat tool from the pycombat package [31], which corrects for artificial differences in the overall expression distribution of each sample by using Location and Scale (L/S) adjustments (**Figure 2B**). We then integrated samples across different diseases or species into a single expression matrix by using the unique gene symbol as key, with genes in rows and samples in columns. The standardized dataset was ready for downstream analysis of differentially expressed genes (DEGs) and for training machine learning models.

**Figure 2.**
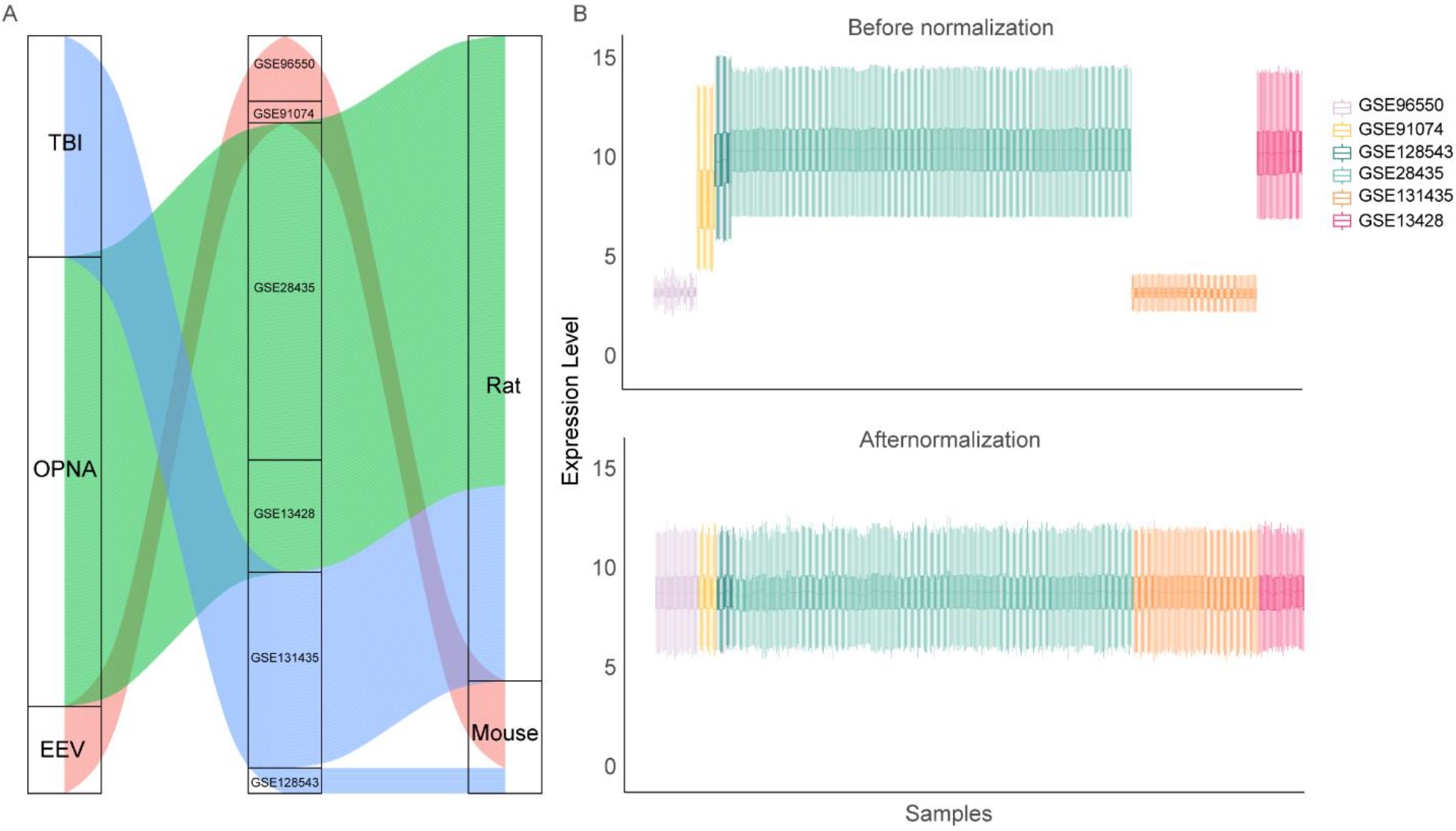
Data collection and normalization in this study. **A)** Public available datasets used in this study across species and diseases. **B)** Gene expression level across multiple datasets before and after combat normalization.

### Differential expression analysis between OPNA/EEV/TBI and control samples in rodent models

To investigate the characteristics of gene expression changes under different conditions, we implemented differential expression analysis between disease and control samples for each dataset separately. For the comparison between EEV/TBI/OPNA and their corresponding normal controls, we identified 2542, 5211, and 766 DEGs, respectively. Interestingly, in all three conditions, there appear to be more up-regulated DEGs than down-regulated ones (**Figure 3A**). To further explore the intrinsic connections of the gene expression changes with respect to these three brain diseases, we crossed the three up- and down-regulated DEGs to determine the common and distinct gene expression changes among different diseases. We found that most DEGs are disease-specific, in other words, the DEGs in all three diseases do not overlap significantly. For up- and down-regulated DEGs, only 45 and 1 DEGs were found to have simultaneous changes in all three diseases, respectively (**Figure 3B**).

**Figure 3.**
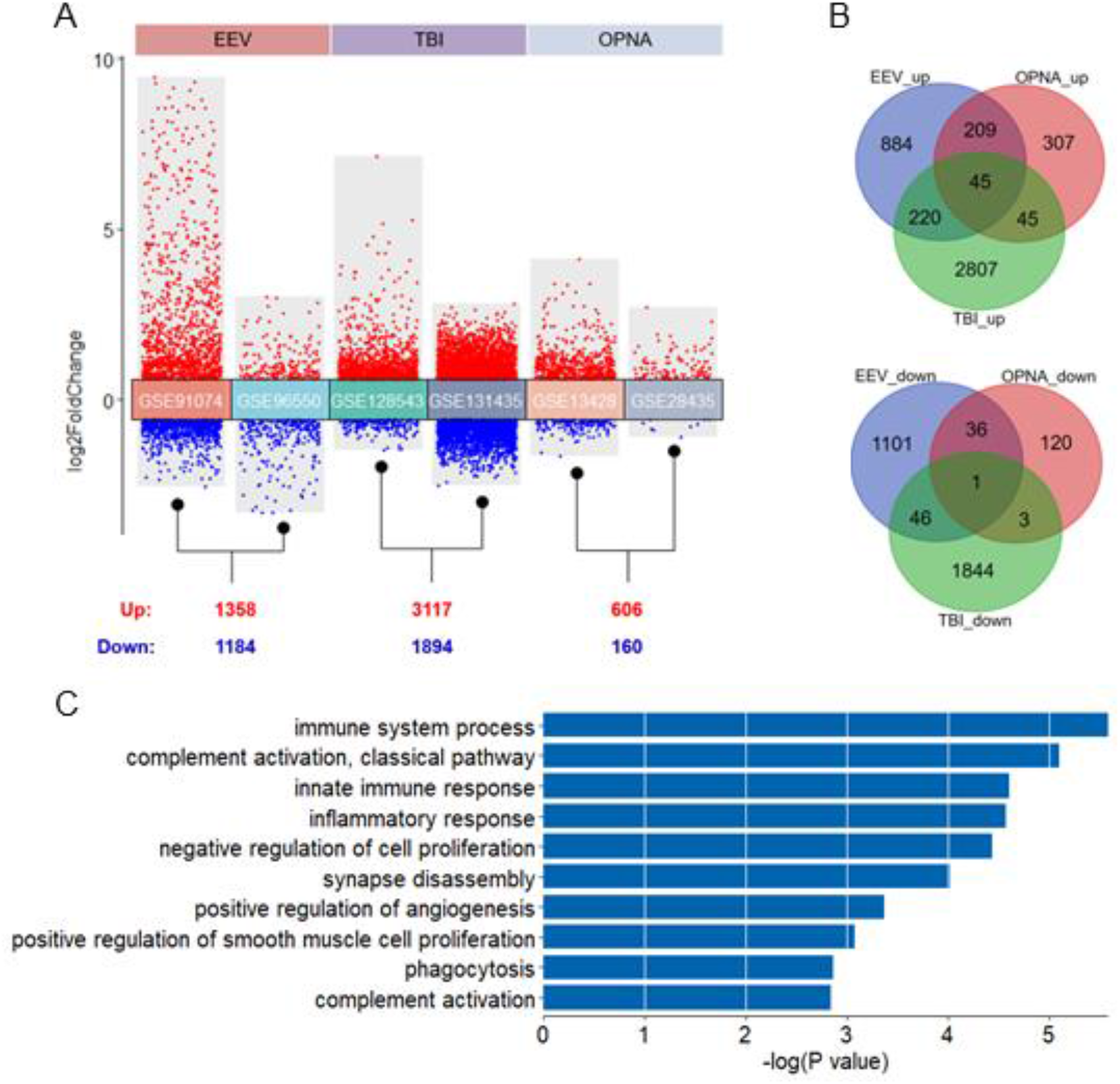
Differential expression analysis between disease and control samples. **A)** Transformed volcano plot depicting differentially expressed genes (DEGs) between disease and control. **B)** Overlap of DEGs among the three types of brain diseases studied. The top panel shows the number of up-regulated DEGs, while the bottom panel displays the down-regulated DEGs. **C)** GO ontology enrichment for 45 DEGs shared in up-regulated DEGs in all three brain diseases.

To analyze the functional consequences resulting from these gene expression changes, gene functional enrichment analysis was conducted on both shared and disease-specific gene sets using the DAVID website [32], by which an over representation analysis was adopted to determine the enriched biological processes of input gene set. As expected, the 45 shared up-regulated DEGs are significantly enriched in immune- and neuron-related functions, such as “immune system process”, “innate immune response”, “inflammatory response”, “synapse disassembly”, and “positive regulation of angiogenesis” (**Figure 3C**). Moreover, we found a list of immune-related and neuron-related processes enriched in disease-specific DEGs, which indicates that specific brain functions may be affected by different brain diseases (**Figure S1**). Above all, our findings provide valuable insights into the molecular mechanisms underlying various brain pathologies and identify potential therapeutic targets for further investigation. The shared gene signature across multiple brain diseases suggests common pathways that could be targeted for broad-spectrum treatments, while the disease-specific signatures offer opportunities for developing targeted therapies for individual conditions.

### Machine learning framework to identify key expression features for OPNA, EEV and TBI

We combined the DEGs identified in each dataset, and from which selected orthologous genes across species to build an informative gene set for machine learning. A total of 2525 such genes were retained for downstream classifier training and prediction. Different machine learning approaches, including k-nearest neighbor (KNN), random forest (RF), linear discriminant analysis (LDA), support vector machine (SVM), and artificial neural network (ANN), were then employed to identify common and unique signatures with respect to OPNA, EEV, and TBI. Under a training to test ratio of 4:1, we obtained the misclassification rate of the five classifiers (**Table 1**), from which we see that the ANN classifier achieves the best performance for both ternary (across the three disorders to identify distinct features for each disorder) and binary (between disease and control to identify common features of three disorders) classification (**Figure 4A&B**). We therefore chose ANN as a suitable machine learning model in this study for its outperformance among others.

**Table 1.**
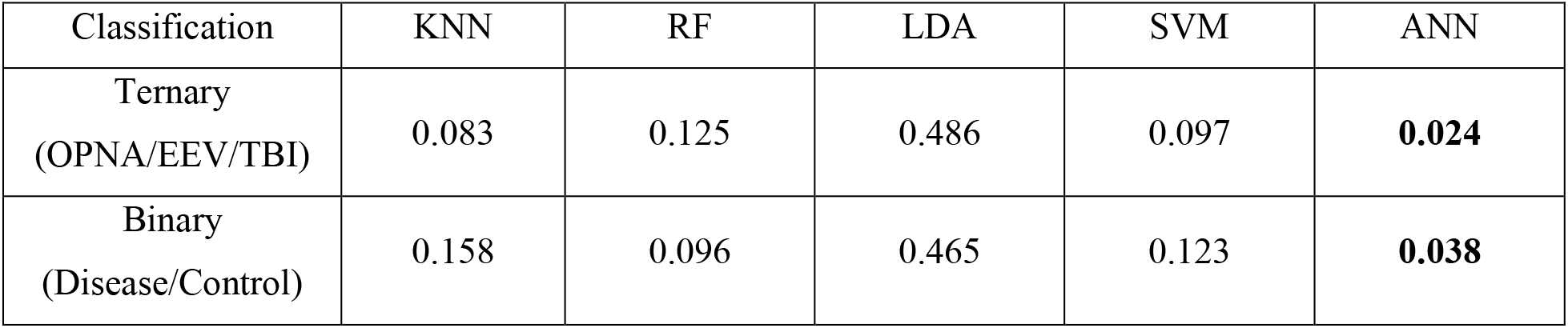
Misclassification rates of different machine learning models.

| Classification | KNN | RF | LDA | SVM | ANN |
| --- | --- | --- | --- | --- | --- |
| Ternary<br>(OPNA/EEV/TBI) | 0.083 | 0.125 | 0.486 | 0.097 | **0.024** |
| Binary<br>(Disease/Control) | 0.158 | 0.096 | 0.465 | 0.123 | **0.038** |

**Figure 4.**
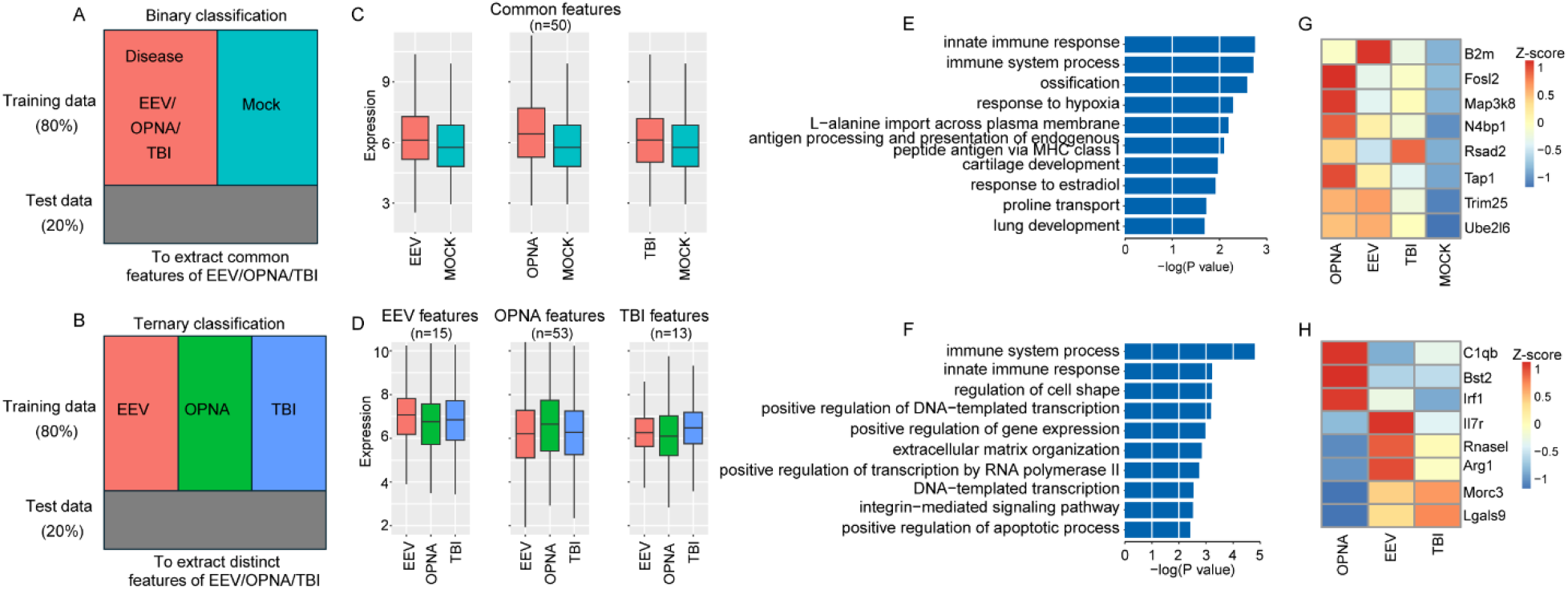
Feature selection results for ANN ternary and binary classification. **A and B)** Sketch Map showing the binary (A) and ternary (B) classification. **C and D)** Average expression of common (C) and distinct (D) expression features. **E and F)** GO ontology enrichment for binary (E) and ternary (F) classification features. **G and H)** Expression of selected examples for immune related features for binary (G) and ternary (H) classification.

We next used the ANN model to select a subset of feature genes that are most important for classification. For ternary and binary classifications, 238 and 252 genes were selected, respectively, and by using these selected feature genes, the ANN classifier was able to achieve the same misclassification rate as by using all genes. To ensure that the selected features are representative for disease in binary classification and for each disorder in ternary classification. We set between group standard variation over 0.15 as an additional threshold to filter the selected features, 50 and 81 features were retained as the final expression features for binary classification and ternary classification. The retained features showed strong expression representative in both binary and ternary classification (**Figure 4C&D**). This provides strong evidence that these selected feature genes do play a critical role in separating the classes, either across OPNA/EEV/TBI or between disease and control.

Lastly, we examined the function of selected feature genes. Functional enrichment analysis showed that both common and distinct features of three brain disorders are significantly enriched in immune-related terms including “immune system process” and “innate immune response” (**Figure 4E&F**). Among these immune-related genes, we identified a core set of upregulated genes across all three disorders compared to mock samples (**Figure 4G**). These included *B2m* (beta-2-microglobulin, crucial for MHC class I antigen presentation), *Fosl2* (a key transcription factor in inflammatory responses), *Map3k8* (a central regulator of inflammatory cytokine production), *Rsad2* (an interferon-stimulated gene with antiviral properties), *Tap1* (involved in antigen processing), *Trim25* (a critical regulator of innate immunity), and *Ube2l6* (involved in ISGylation) were consistently upregulated, suggesting a common inflammatory signature across these conditions. Among the ternary classification features, we identified disorder-specific immune signatures (**Figure 4H**). OPNA samples showed elevated expression of *C1qb* (complement cascade initiator), *Bst2* (type I interferon-induced antiviral protein), and *Irf1* (interferon regulatory factor). EEV specifically upregulated *Il7r* (lymphocyte development regulator), *Rnasel* (viral RNA degradation), and *Arg1* (immunosuppressive mediator in myeloid cells). TBI samples distinctively expressed *Morc3* (nuclear protein involved in immune response) and *Lgals9* (immunomodulatory galectin). Above all, through machine learning algorithms, we have successfully extracted the common and distinctive expression features underlying EEV, TBI, and OPNA.

## Conclusions and discussion

In this study, we developed and implemented a comprehensive framework for analyzing and comparing transcriptomic profiles across three distinct neurological conditions: TBI, EEV, and OPNA. By leveraging machine learning approaches, particularly artificial neural networks, we identified both shared and condition-specific gene signatures that provide valuable insights into the underlying molecular mechanisms of these neurological disorders. Our comparative analysis revealed several key findings. First, the integration of diverse transcriptomic datasets demonstrated the feasibility of cross-platform data normalization and analysis. Second, the machine learning models enabled identification of critical genes associated with each condition, suggesting potential therapeutic targets for medical countermeasures. Third, the pathway and gene ontology analyses highlighted specific biological processes and molecular functions that may play crucial roles in the pathogenesis of these neurological conditions.

Several limitations exist in our study. First, OPNA and EEV studies are highly limited, therefore limiting our ability to have consistent tissue types in the data selected. Future studies would benefit from incorporating additional neurological conditions with more abundant data from the same species, tissue type, and background.. Next, our analysis relies on microarray data, which provides only tissue-level expression profiles and may overlook cell type-specific responses that are crucial for understanding the complex pathophysiology of neurological disorders. Additionally, microarray technology has inherent limitations in detecting novel transcripts and may identify fewer transcripts compared to more recent sequencing approaches. Future studies could address these limitations by incorporating single-cell RNA sequencing (scRNA-seq) data to identify cell type-specific responses to TBI, EEV infection and OPNA exposure and reveal cellular heterogeneity within affected tissues.

Regarding the specific findings of this study, the overwhelming similarities between EEVs, TBIs, and OPNAs are associated with the upregulation of the immune response. While this is an expected finding, it is also challenging to utilize broad immune markers for biomarker analysis or therapeutics. Binary analysis of neurological disease phenotypes could be broadened to include other neuropathologies to identify signatures of gene expression which are indicators of damage in cases where the injury is unknown. Especially in the context of the war fighter, early markers of inflammation could be beneficial in distinguishing a healthy phenotype, where immune associated genes are lowly expressed, from a recent injury or exposure to chemical or biological agents where these genes are upregulated. Further research is required to assess whether these biomarkers of disease in the brain, also correlate with differences in the blood to make sample collection and testing realistic in the field. For the tertiary analysis (EEV vs TBI vs OPNA), these results could be utilized to characterize types of injury. For example, in this analysis the gene Morc3 or Lgals9 appear to be upregulated in OPNA exposure and downregulated in EEVs. Therefore, these genes could be a feasible biomarker in clinical setting to distinguish whether an individual posing general malaise like symptoms could have been exposed to a viral or chemical threat. This method of biomarker analysis has previously been used in the clinical context to assess whether patient inflammation is associated with infection, where supportive antimicrobial therapeutics are necessary, or an underlying disease state such as cancer, ischemia, or pulmonary embolism, where administration of antimicrobial agents could worsen disease and in some cases be fatal [33]. As there are few comparable transcriptomic studies for EEVs and OPNA. Further validations and incorporation of additional datasets is crucial for assessing the feasibility of transcriptional biomarker identification.

Our findings provide a prospect for future investigations into these neurological conditions and demonstrate the value of machine learning in understanding complex disease mechanisms. As we move forward with single-cell approaches, we expect to gain even deeper insights into the cellular and molecular mechanisms underlying these disorders, ultimately contributing to the development of more effective medical countermeasures.

## Methods

### Data collection, normalization, and integration

The publicly available gene expression data were obtained from the National Center for Biotechnology Information (NCBI), under the Gene Expression Omnibus (GEO) database. A total of 395 samples were used in this study, for investigating the transcriptional responses of different diseases such as EEV, TBI, and OPNA exposure. These datasets were obtained from various experiments across mammalian species, primarily rats and mice, summarized in **Table 2**. Rodent data for TBI, OPNA, and EEV were selected based on data availability and an attempt to limit to areas of the brain which could be most correlative but is also limited because of minimal data for EEVs and OPNAs. Each dataset was filtered to remove knockouts, treatments, or additional treatments to ensure these data did not impact results (**Table S1**).

**Table 2.** Resource of datasets used in this study.

| Data | Data Type | Disease Type | Brain Location | Species | Title | PMID |
| --- | --- | --- | --- | --- | --- | --- |
| GSE91074 | Microarray | EEV | Whole Brain | Mouse | Gene expression in the brains of different strains of laboratory mice upon intranasal infection with vaccine strain (TC83) of Venezuelan equine encephalitis virus | 28184218 |
| GSE96550 | Microarray | EEV | Whole Brain | Mouse | Differential host gene responses from infection with neurovirulent and partially-neurovirulent strains of Venezuelan equine encephalitis virus | 28446152 |
| GSE128543 | Microarray | TBI | Cortex | Mouse | Identification of Novel Targets of RBM5 in the Healthy and Injured Brain | 32335213 |
| GSE131435 | Microarray | TBI | Hippocampus | Rat | TBI weight-drop model with variable impact heights differentially perturbs hippocampus-cerebellum specific transcriptomic profile. | 33172833 |
| GSE13428 | Microarray | OPNA | Hippocampus | Rat | Gene Expression Profiling of Rat Hippocampus Following Exposure to the Acetylcholinesterase Inhibitor Soman | 19281266 |
| GSE28435 | Microarray | OPNA | Piriform cortex | Rat | Transcriptomic Analysis of Rat Brain Following Exposure to the Organophosphonate Anticholinesterase Sarin | 21777429 |

All the samples were integrated into one expression matrix with genes listed in rows and samples in columns, by merging genes with the same symbol in different samples or species. ComBat [31] was adopted to normalize the data across platforms and experiments and adjust for batch effects. The normalized expression matrix, with dimension 6289×395, was retained and used for downstream analysis.

### Differential expression analysis

The Limma package [34] in R was adopted to identify differentially expressed genes (DEGs) between the disease samples and control samples for EEV, OPNA, and TBI, respectively. An empirical-Bayes based method was used to determine the expression difference and statical significance, so that genes with fold-change over 1.5 and p-value less than 0.05 were identified as DEGs.

### Gene functional enrichment analysis

Gene functional enrichment analysis was conducted using the DAVID website [32], which implemented a hypergeometric test to evaluate the enrichment score, and the significance of the input gene sets in certain gene ontology (GO) terms. The GO terms with p-value less than 0.05 were determined as the over-represented biological processes.

### Machine learning on transcriptomic datasets

Machine learning models, particularly deep learning ANNs, were used to predict brain disease based on the normalized expression matrix. To enhance model efficiency, only DEGs obtained by contrasting the disease samples and control samples were included in the models, reducing the dimension from 6289 to 2525. Both ternary and binary classifications were implemented to compare prediction accuracy across various conditions, where the former used OPNA, EEV, and TBI as responses and the latter used disease (by combining samples from three disorders) and control (by combining corresponding control samples) as responses. Specifically, for KNN classification, we set *k* = 3; for RF classification, we used 1000 decision tree; and for SVM classification, we chose radial basis kernel. For ANN classification, the entire dataset was split into training and testing sets, according to a ratio of 4:1. The ANN model consists of three hidden layers, with the number of nodes in each layer specified according to the geometric pyramid rule [35]. Model performance was subsequently evaluated using the misclassification rate.

Besides prediction, ANNs were also used to identify feature genes that are shared or unique among different disorders. Such a feature selection was achieved by heuristically searching the 2525-dimensional feature space along certain path to find the most parsimonious model that attains comparable or slightly higher misclassification rate (MCR) to the full model. The search path was determined by calculating the leave-one-out model MCR and ranking the marginal effect of each gene.

## Acknowledgments

This project was supported by the Defense Threat Reduction Agency (HDTRA1-23-1-0009).

## Author contribution statement

X. W., K. K. and H. X. conceived the experimental design. L. Y., M. V., V. K., B. C., P. C., and X. W. collected and analyzed the data. L. Y., M. T., E. J., K. K., X. W., and H X interpreted the results and drafted the manuscript. All authors discussed the results, read, and edited the manuscript, and approved the final manuscript.

