## Supplementary figures and images for "Machine Learning Identifies Genes Linked to Neurological Disorders Induced by Equine Encephalitis viruses (EEV), Traumatic Brain Injuries (TBI), and Organophosphorus nerve agents (OPNA)"

### Supplemental Figure 1

#### Figure S1

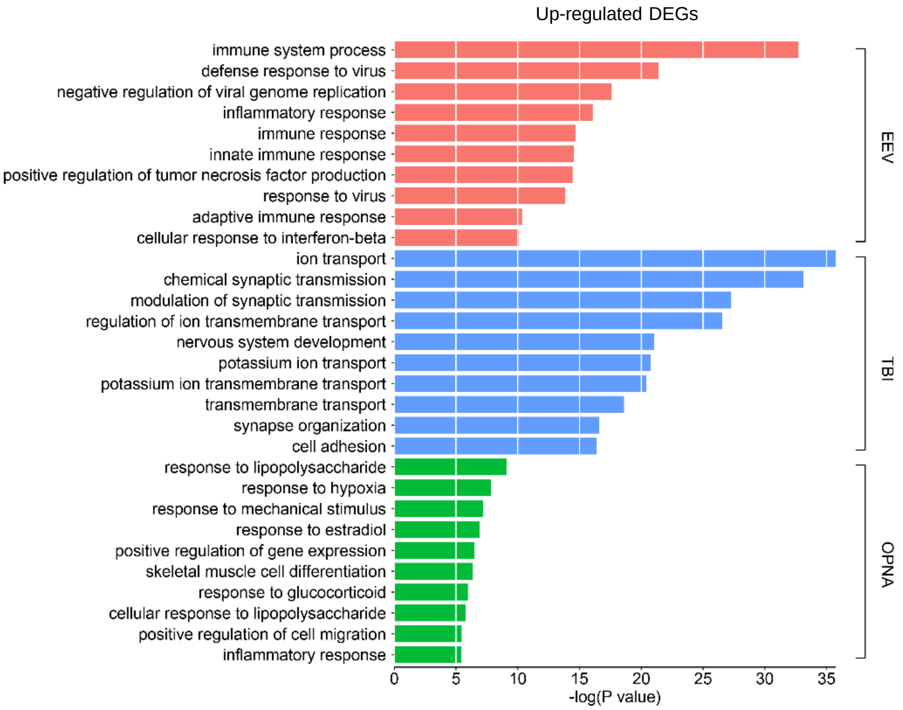


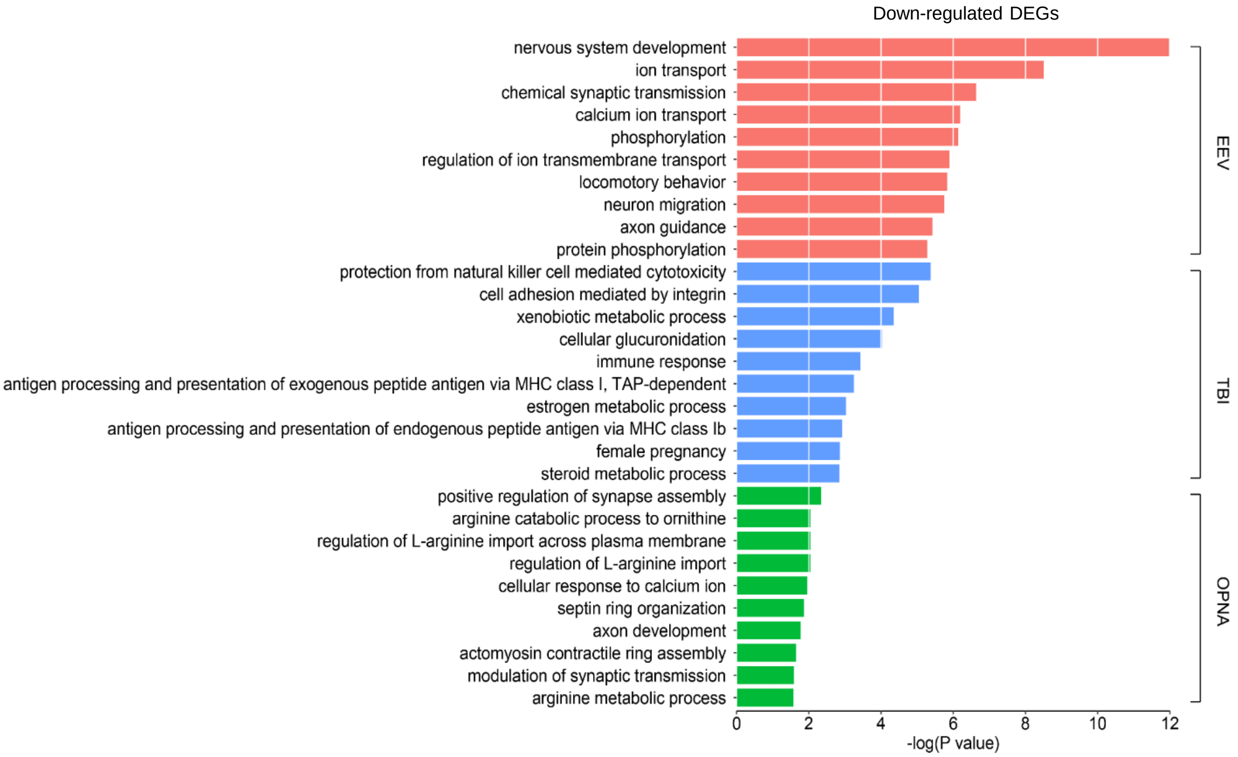


**Figure S1. GO ontology enrichment for disease-specific up- and down-regulated DEGs.**
